# Binocular Listing’s Law For Human Eyes’ Misaligned Optical Components

**DOI:** 10.1101/2024.06.24.600541

**Authors:** Jacek Turski

## Abstract

Human eyes’ optical components are misaligned. This study presents the geometric construction of ocular torsion in the binocular system, in which the eye model incorporates the fovea that is not located on and the lens that is tilted away from the eye’s optical axis. The ocular torsion computation involves Euler’s rotation theorem in the framework of Rodrigue’s vector. When the eyes’ binocular posture changes, each eye’s torsional orientation transformations are visualized in *GeoGebra’s* dynamic geometry environment. Listing’s law, which originally restricts single-eye torsional positions and has imprecise binocular extensions, is formulated *ab initio* for binocular fixations using Euler’s rotation theorem. It replaces Listing’s plane and related perpendicular primary direction with the empirical horopter’s abathic distance fixation, which has a straight frontal line configuration at this fixation. Notably, it corresponds to the eye muscles’ natural tonus resting position, which serves as a zero-reference level for convergence effort, providing the missing neurophysiological significance of Listing’s plane. Thus, Listing’s law use in clinical diagnosis and management of strabismus should be updated.

## 1 Introduction

In healthy human eyes, the fovea is not located on, and the crystalline lens is tilted away from the eye’s optical axis. This asymmetry has been studied in many clinical papers, usually decomposed along temporal-nasal and inferior-superior axes, that is, into horizontal and vertical segments, respectively [Chang et al., 2007, de Castro et al., 2007, Schaeffel and Kaymak, 2010, Aguirre, 2019, Wang et al., 2019]. The fovea’s anatomical displacement, relatively stable in the human population Holladay [2007], and the cornea’s asphericity contribute to optical aberrations, and the lens tilt partially compensates for these aberrations [Tabernero et al., 2007, Charman and Atchison, 2009, Artal, 2014, Liu and Thibos, 2017].

Recently, the geometric theory of the binocular system with the asymmetric eyes (AE) that comprises the fovea’s horizontal displacement from the posterior pole and the lens horizontal tilt relative to the eye’s optical axis has been developed in a series of papers Turski [2018, 2020, 2023]. Using this theory, the impact of the eye’s misalignment on stereopsis organized by the horizontal retinal corresponding points—for an element in the retina of one eye, there is the corresponding element in the other eye that shares one visual direction— was finally explained.

The AE with vertical misalignments of optical components, complementing the considered before horizontal misalignment, was introduced in Turski [2024]. The impact of these misalignments on binocular vision is briefly discussed in the next section, which should help read the rest of the paper.

The study presented here starts with the geometric definition of ocular torsion in the AE about the lens’s optical axis direction, which best approximates the usually considered ocular torsion about the visual axis of the axially symmetric schematic eye. This geometric ocular torsion is implemented in the binocular system for each AE, and the torsional disparity is calculated for the 3D binocular field of fixations. The tools used in the theory consist of Euler’s rotation theorem in Rodrigues’ (rotation) vector framework and visualization in *GeoGebra’s* dynamic geometry simulations. It provides a consistent binocular formulation of Listing’s law for the first time.

Crucially, the Listing plane and the primary direction used in constructing Listing’s law must be replaced with the eyes’ binocular resting posture corresponding to the abathic distance fixation of the empirical horopter distinguished by its straight-line shape and the frontal orientation. Thus, the results of this study indicate that Listing’s law should be updated to address the control of 3D eye movements, clinical implications for strabismus, and optimal management of this disorder.

## 2 Prior Studies with the AE

In the eye model introduced in Turski [2018], the lens is represented by (i) the nodal point located on the optical axis 0.6 cm anterior to the eye’s rotation center and (ii) the image plane parallel to the lens’s equatorial plane and passing through the eye’s rotation center. In the latest paper Turski [2024], based on the clinical studies listed in Section 1, asymmetry angles for the fovea’s horizontal and vertical displacements from the posterior pole are *α* = 5.2^*°*^ and *γ* = −2^*°*^ and the lens’ horizontal and vertical tilts relative to the optical axis are *β* = 3.3^*°*^ and *ε* = −1^*°*^, explained in Figure 1 for the right AE. The left AE is the mirror-symmetric for eyes fixating in the median plane. In Figure 1, the axis *H*_*r*_ is the image plane axis of retinal horizontal correspondence—the location of the corresponding points used to construct the iso-disparity curves—the retinal disparity spatial coordinates. The image plane axis 𝒱_*r*_, which is parallel to the lens’s vertical direction in its equatorial plane (the green line through *N*_*r*_), is the location of the points of retinal vertical correspondence used to construct the vertical horopter. This axis is tilted by *η* = 0.183^*°*^ relative to the true vertical axis *V*_*r*_ with its top in the temporal direction.

**Figure 1:**
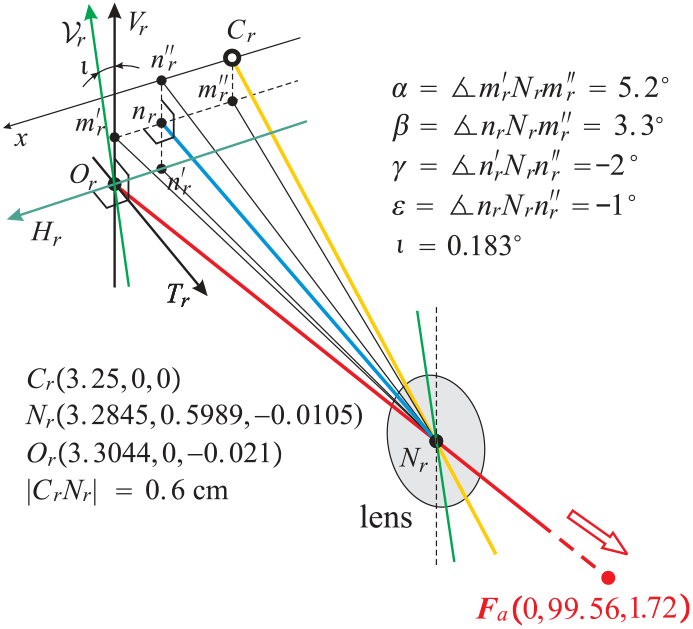
The right 3D AE near the optical center *O*_*r*_, the eye’s rotation center *Cr* and the nodal point *N*_*r*_. The yellow line is the eye’s optical axis, the red line is the visual axis fixating *F*_*a*_, and the blue line is the lens’s optical axis. The AE’s image plane coordinates consist of the axis *H*_*r*_ of the intersection by the visual axis and the vertical axis *V*_*r*_, both attached at *O*_*r*_. They are complemented by the perpendicular coordinate *T*_*r*_ parallel to the lens’s optical axis. The coordinate axes *H*_*r*_ and *𝒱*_*r*_ express the horizontal and vertical correspondence.

The geometric theory of the longitudinal horopter in the binocular system with the 2D AE comprising only horizontal misalignment of the fovea and lens was developed in Turski [2020]. It was further extended to iso-disparity conic sections and applied in the study of stereopsis for the horizontal fixations in Turski [2023]. I recall that stereopsis—the perception of objects’ form and their location in 3D space Wheatstone [1838], Julesz [1971]—is organized by the horizontal retinal correspondence: For a small retinal area in one eye, there is a corresponding unique area in the other eye such that both share one subjective visual direction. Finally, in Turski [2024], 3D AE with the horizontal and vertical misalignment angles mentioned above and the 3D binocular field of fixations were used to study stereopsis and the subjective vertical horopter.

The binocular system with the AEs has a unique fixation for the given asymmetry angles *α, β, γ, ε* such that the resulting posture consists of the vertical coplanar image planes parallel to the coplanar lenses’ equatorial planes. The simulations in *GeoGebra’s* dynamic geometry environment demonstrated in Turski [2023, 2024] that in this posture, the iso-disparity curves are frontal straight lines. Importantly, it corresponds to the eye muscles’ natural tonus resting position, which serves as a zero-reference level for convergence effort [Ebenholtz, 2001], providing the missing neurophysiological significance of the primary position (Listing’s plane and primary direction). As emphasized by Hess and Thomassen [2014], the neurophysiological significance of the primary position has remained elusive despite its theoretical importance in oculomotor research.

For the values of asymmetry angles listed above, the fixation of the special posture is *F*_*a*_(0, 99.56, 1.72) with coordinates in centimeters, corresponding to the average abathic distance fixation of about 1 m. Importantly, this binocular posture, which turns out to be crucial in the new formulation of Listing’s law, is referred to in Turski [2023, 2024] as the eyes’ resting posture (ERP) because the average asymmetry angles of the AE numerically correspond to the eyes’ resting vergence posture corresponding to the eye muscles’ natural tonus resting position mentioned above.

The main contributions resulting from the studies mentioned above are as follows:

1. The asymmetric retinal correspondence—the corresponding elements are compressed in the temporal retinae relative to those in the nasal retinae—which is person-dependent and difficult to measure [Shipley and Rawlings, 1970], is represented by the symmetric distribution in the image plane of the ERP by AE’s horizontal misalignment angles. The correspondence symmetric distribution on the image plane is preserved for all fixations [Turski, 2020].
2. The asymmetry of the retinal correspondence is caused by the horizontal misalignment of the fovea and the lens in healthy human eyes. It is demonstrated in Turski [2023] by the universality of the spatial iso-disparity distribution for the AE’s angles of horizontal misalignment. This means that the iso-disparity line distribution is invariant to all *β* values in the AE.
3. The Helmholtz’s vertical retinal criterion of the tilted vertical meridian, which is less relevant for misaligned eye’s optical components modeled with the AE, is replaced by the tilted lens’ vertical retinal criterion [Turski, 2024]. This new criterion provides the experimentally observed inclination of the vertical horopter that agrees with results in Amigo [1974], Siderov et al. [1999], in contrast to the values provided by Helmholtz [1867/1925] and others.

In Turski [2023], the iso-disparity curves consist of families of ellipses or hyperbolas such that the zero-disparity horopters resemble empirical horopters. Moreover, in the framework of Riemannian geometry, this reference studied the global aspects of phenomenal spatial relations, such as visual space variable curvature and finite horizon. The global aspects of stereopsis are crucial in perception, as exemplified by the coarse disparity contribution to our impression of being immersed in the ambient environment despite receiving 2D retinal projections of the light beams reflected by spatial objects [Barry, 2013].

## 3 Method: *GeoGebra* Simulations and Geometric Constructions

### 3.1 *GeoGebra* Simulations

*GeoGebra* simulations of the eyes’ binocular postures include geometric primitives (lines, conics, planes, spheres, cylinders, etc.), geometric constructs (tangent, bisector, etc.), and basic geometric transformations (translations, rotations, etc.). These geometric tools are used directly to design the visualization, so programming is not involved. However, the main problem for creating a dynamic simulation in this study is to connect those geometric tools such that by moving the fixation point, the binocular system with AEs accordingly changes, visualizing its transformations. In the simulations presented in this study, the number of geometric tools is quickly growing, and those that create connections are hidden.

Because the *GeoGebra* does not involve programming, the links to simulations of ocular torsion in the binocular system with AEs are available upon reasonable request.

### 3.2 Ocular Torsion and the AE

Ocular torsion is usually described as a rotation in the eyeball around the line of gaze. This definition is impractical for the AE because it includes misaligned optical components. The geometrical ocular torsion of the AE is defined as a rotation around the image plane perpendicular coordinate axis at the optical center—the axis *T*_*r*_ for the right AE in Figure 1. Because this axis is parallel to the lens’s optical axis at a distance of 0.022 cm, it gives the angle of 2^*°*^ between the axis *T*_*r*_ and the visual axis (the red line) in Figure 1.

How the ocular torsion is computed and simulated in *GeoGebra* is explained with the help of Figure 2 and discussed later for the actual simulations in Section 4.

**Figure 2:**
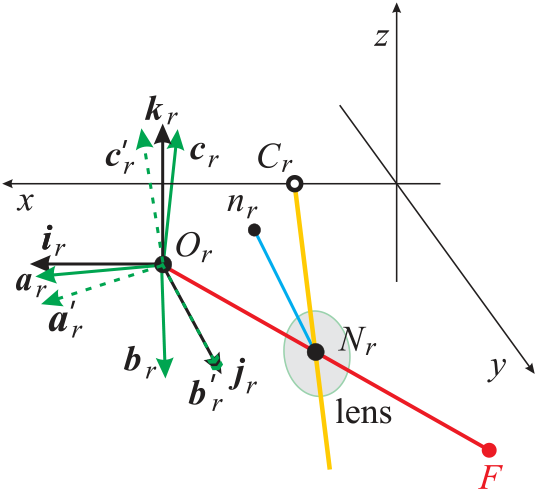
Schematic explanation of how the ocular torsion is geometrically defined and calculated for the right eye. The black frame at *O*_*r*_ agrees with the axes *H*_*r*_, *T*_*r*_, *V*_*r*_ in Figure 1 and with the head coordinates *x, y, z*. The green frame is the transformed black frame after the posture changed from the fixation at *F*_*a*_ in the ERP to a fixation *F*. The green ‘primed’ frame in dashed lines shows the rotated green frame in the plane containing the vectors **b***r* and **j**_*r*_ that overlays **b**_*r*_ with **j**_*r*_. The geometric definition of the ocular torsion is the angle 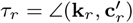.

By the definition of ocular torsion given in Figure 2, the rotation vector of the black frame is contained in the image plane of the ERP, and ocular torsion is about the line through *O*_*r*_ parallel to the lens’s optical axis (the line through *n*_*r*_ and *N*_*r*_) which is perpendicular to the image plane. This result establishes the geometrical calculations of the eyes’ ocular torsions for any binocular fixation executed from the ERP. The binocular ERP provides a neurophysiologically motivated new definition for Listing’s law eye’s primary position originally considered for the single eye; further discussed in Section 5.

### 3.3 Examples: Simulations of Iso-disparity conics sections and Vertical Horopter

In this section, I visualize in *GeoGebra* the longitudinal disparity spatial coordinates (iso-disparity curves) and the subjective vertical horopter in the ERP of fixation *F*_*a*_(0, 99.56, 1.72) and the tertiary posture after the fixation point *F*_*a*_ changed to *F* (12, 33.56, 5.72). They are shown in Figure 3. The details of these and other simulations are presented in Turski [2023].

**Figure 3:**
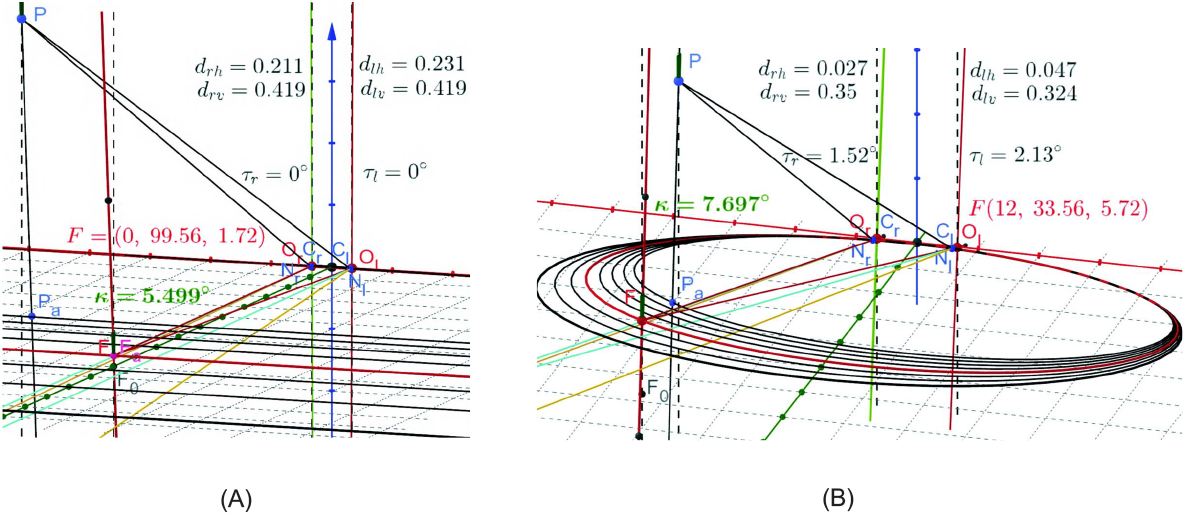
(A) The ERP with the fixation at *Fa* showing the iso-disparity frontal lines in the visual plane and the vertical horopter tilted by *κ* = 5.5^*°*^ relative to the objective vertical. The longitudinal and vertical horopters passing through *F*_*a*_ are shown in red. The point *F*_0_ is the projection of *F*_*a*_ into the head’s transverse plane containing the eyes’ rotation centers (marked with the grid). (B) The simulation of the iso-disparity ellipses and the vertical horopter after the fixation at *F*_*a*_(0, 99.56, 1.72) is changed to *F* (12, 33.56, 5.72). The vertical horopter is tilted by *κ* = 7.7^*°*^ in the plane 1.3*x* + 0.12*y* = 20. The calculated disparities are discussed in the text.

I should clarify that the iso-disparity curves and the subjective vertical horopter are precisely defined in the binocular system with AEs only for fixations in the median plane [Turski, 2024]. The isodisparity conic sections in Figure 3 (A) are straight frontal lines in the ERP. Also, in this posture, the vertical horopter is a straight line tilted from the true vertical by *κ* = 5.5^*°*^ with its top away from the head. From the data in Figure 3 (A), the horizontal disparity in coordinates (*H*_*r*_, 𝒱_*r*_) for the right AE and (*H*_*l*_, 𝒱_*l*_) for the left AE given by *δ*(*P*) = − 0.231 0.211 cm. The vertical disparity and the torsional disparity are zero.

Figure 1 (B) shows the iso-disparity ellipses and the vertical horopter for the tertiary fixation *F* (12, 33.56, 5.72). The horizontal disparity value is *δ*(*P*)_*h*_ = 0.02 cm, the vertical disparity value is *δ*(*P*)_*v*_ = 0.026 cm and the torsional disparity value is *δ*_*t*_ = *τ*_*r*_− *τ*_*l*_ =−0.61^*°*^.

The projection of *P* into the visual plane along the parallel line to the vertical horopter, *P*_*a*_, is exactly on the fourth consecutive disparity line down from the horopter (red frontal line through *F*_*a*_). Because the horizontal retinal relative disparity value between each consecutive disparity line is 0.005 cm for the simulations, the spatial disparity of *P* agrees with the retinal horizontal disparity in both panels (A) and (B) of Figure 3,

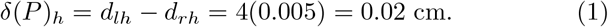

I only mention that this result indicates that the subjective vertical horopter agrees with the perceived vertical, and I refer the reader to Turski [2024] for a detailed discussion of the simulated cases in Figure 3 and other important cases.

## 4 Simulation of Ocular Torsion

The actual geometric construction in *GeoGebra’s* simulation shown in Figure 4 computes each of the two eyes’ ocular torsion for the case in Figure 3 (B) with the fixation point denoted here by *F* ^1^. In Figure 4, the solid green frame for the right AE, (**a**_*r*_, **b**_*r*_, **c**_*r*_), and the brown frame for the left AE, (**a**_*l*_, **b**_*l*_, **c**_*l*_), are moving with the image planes from their position that initially in ERP agrees with the respective black frame for each AE.

**Figure 4:**
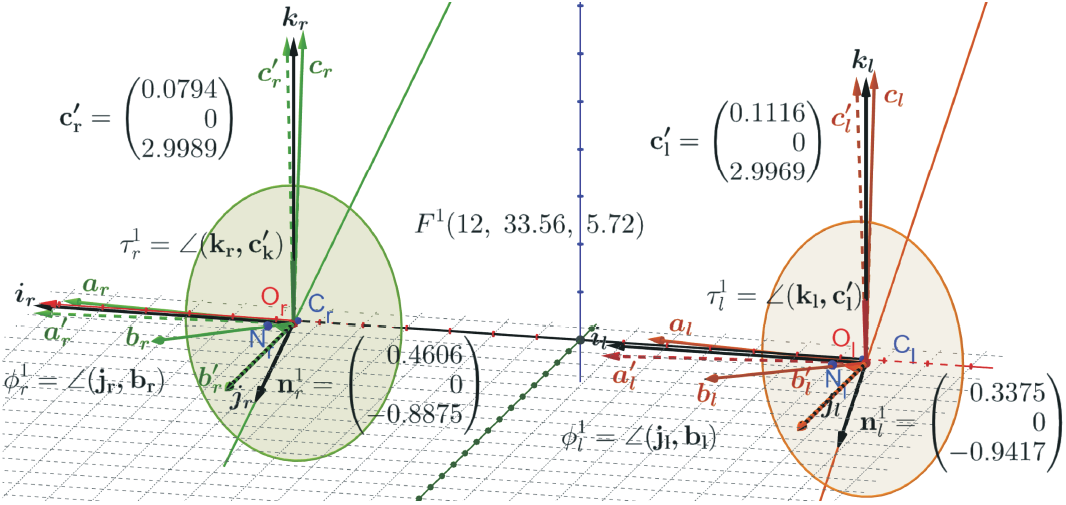
The black frames in the ERP (**i**_*r*_, **j**_*r*_, **k**_*r*_) and (**i**_*l*_, **j**_*l*_, **k**_*l*_) for the right and left AEs are attached at the image planes optical centers *O*_*r*_ and *O*_*l*_. Their orientations agree with the upright head’s frame. The solid-colored frames are moving from the initial position of black frames for a given fixation *F* ^1^. The dashed-colored frames are obtained by rotating the solid-colored frames by moving **b**s vectors onto the **j**s vectors. The rotation axes are given by the vector 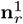 and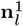, with the angle of rotations 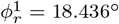 and 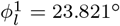, are contained in the co-planar image planes of the ERP, parallel to the head frontal plane. The ocular torsion of each AE is 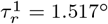 and 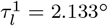.

To calculate each eye’s ocular torsion, the green and brown frames are rotated such that **b**s vectors are overlaid with **j**s vectors. The rotations are in the planes spanned by **b**s and **j**s vectors such that rotation axes containing the vectors 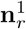 and 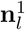 are perpendicular to these planes. The rotations result in two ‘primed’ frames shown in dashed lines. The rotations angles, ∠(**j**_*r*_, **b**_*r*_) = 18.436^*°*^ and ∠(**j**_*l*_, **b**_*l*_) = 23.821^*°*^ are denoted by 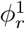 and 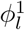.

Because 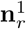 is perpendicular to **j**_*r*_ and 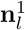 is perpendicular to **j**_*l*_; they are in the coplanar image planes for the ERP. The ocular torsion, defined by the angles ∠(**k**_*r*_, **c**^*′*^_*r*_) = 1.517^*°*^ and ∠(**k**_*l*_, **c**^*′*^_*l*_) = 2.133^*°*^, are denoted by *τ* ^1^ and *τ* ^1^ such that the torsional disparity 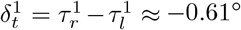. All counterclockwise right-hand rotations are positive.

The results establish the geometrical calculations of the eyes’ ocular torsions for any binocular fixation executed from the ERP. The binocular ERP provides a neurophysiologically motivated new definition for the eye’s primary position, originally part of Listing’s Law that controls ocular torsion but has been considered for the single eye.

## 5 Discussion of Listing’s Law in the Binocular System with the AEs

### 5.1 Ocular Torsion and Listing’s Law

Historically, the torsion of the eye was determined by Donders, Listing, and Helmholtz between 1848 and 1867. Donders’s law is a general statement for a single eye with the head erect and looking at infinity; any gaze direction has a unique torsional angle, regardless of the path the eye follows to get there. Listing’s law, as formulated by Helmholtz in 1867, restricts Donders’s law by stating that visual directions of a single eyesight are related to the eye rotations so that from a reference position, all rotation axes lie in a plane, today referred to as the displacement plane. When the reference direction is perpendicular to the displacement plane, the plane is referred to as the Listing plane, and the unique reference direction is the primary direction. The displacement plane for a given eye’s reference position was studied in the quaternion framework in Tweed et al. [1990], Tweed and Vilis [1990] and reviewed in Haslwanter [1995]. It was only recently numerically simulated in Novelia and O’Reilly [2015].

Listing’s law has been extended in a few ways. In the first extension, Listing’s plane must tilt by half the angle of eye eccentricity from the primary direction Mok et al. [1992], Minken and van Gisbergen [1994]. This monocular rule is known as the *half-angle* rule. The next two extensions involve the two eyes. First, Listing’s law is extended to “binocular” conjugate eye rotations with parallel gazes. Second, for the eyes’ posture with converging gazes on a near fixation point, each eye’s displacement plane turns by the amount proportional to the initial value of vergence angle Van Rijn and Van der Berg [1993], Tweed [1997]. The extensions are known as binocular Listing’s law (L2), whereas the original law is L1.

However, the proportionality coefficients are controversial Van Rijn and Van der Berg [1993]. Further, the primary position, which is still a part of L2 law, corresponds to the eyes’ movements with parallel gazes, and, consequently, its neurophysiological significance has remained elusive despite its theoretical importance in the oculomotor research Hess and Thomassen [2014].

### 5.2 Listing’s Law and the AE: Rodriques’ Vector Computational Framework

Foremostly, the AEs’ image planes for the ERP binocular posture are coplanar and constitute the frontal plane of the stationary up-right head. The gaze direction of each eye in this posture passes through the point *F*_*a*_(0, 99.56, 1.72), expressed in centimeters, and is not perpendicular to this plane, so it is not Listing’s plane. It is not a displacement plane because its orientation is uniquely given by the AE’s parameters *α, β, γ*, and *ε*.

The geometric theory developed here for binocular fixations is based on Euler’s rotation theorem. This theorem states that any two orientations of a rigid body with one of its points fixed differ by a rotation about an axis specified by a unit vector passing through the fixed point. I recall that fixed-axis rotations are curves of the shortest length (geodesics) on the manifold of the rotation (Lie) group *SO*(3) and, hence, are *optimal* ; see Novelia and O’Reilly [2015] for a discussion of *SO*(3) geodesics in the context of eye rotations.

Euler’s rotation theorem is used here in the framework of Ro-drigues’ vector for each of the two eyes constrained by their bifoveal fixations because it can be directly applied in simulations to rotate geometric objects in *GeoGebra’s* dynamic geometry environment. To introduce this framework, I start with the conclusion from Euler’s rotation theorem: Any rotation matrix *R* can be parametrized as *R*(*ϕ*, **n**) for a rotation angle *ϕ* around the axis with the unit vector **n**. This parametrization is unique if the orientation of 0 *< ϕ <* 180^*°*^ is fixed. Usually, a counterclockwise (or right-hand) orientation is chosen to get angles’ positive values.

The pair *ρ* = cos(*ϕ/*2), **e** = sin(*ϕ/*2)**n** is known as *Euler-Rodrigues parameters* and

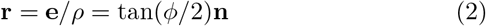

as *Rodrigues’ vector*, usually referred to as *rotation vector* Piña [2011]. Rodrigues proved that under composition *R* = *R*_2_*R*_1_, the corresponding Euler-Rodrigues parameters transform as follows Gray [1980],

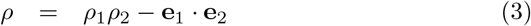

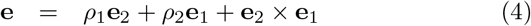

Then, it is easy to see that the corresponding composition for Rodrigues’ vectors is

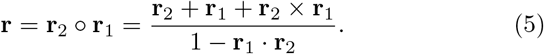

Further, **r**^*−*1^ = − **r** and tan(− *ϕ/*2)**n** = tan(*ϕ/*2)(− **n**) define the same rotation vector.

Note that (cos(*ϕ/*2), sin(*ϕ/*2)**n**) is a unit quaternion that describes the rotation by *ϕ* around **n**, and Rodrigues used in 1840 the multiplication of quaternions to obtain Eqs (3) and (4). Remarkably, the quaternions were formally defined in 1843 by Hamilton, and vectors appeared late in the 19th century when Gibbs and Heaviside independently developed vector algebra.

The rotation vectors in the simulation shown in Figure 4 for fixation *F* ^1^(12, 33.56, 5.72) are

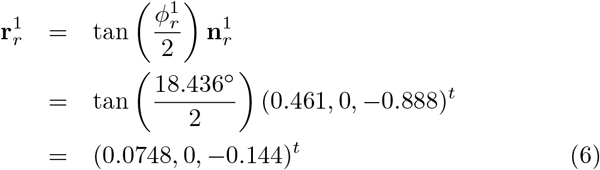

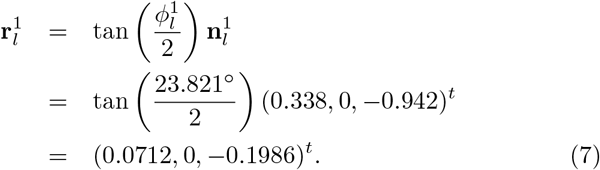

for the right and left eye, respectively.

Now, I can compute directly the torsion changes between tertiary eyes’ binocular fixations, which usually are obtained in oculomotor research using Listing’s L2.

To this end, I first show in Figure 5 the simulation for the change of the fixation points from the resting eyes’ posture at *F*_*a*_(0, 99.56, 1.72) to *F* ^2^(7, 53.56, 15.72). Following similar calculations for *F* ^2^ as before for the fixation *F* ^1^, from results presented in Figure 5, I obtain,

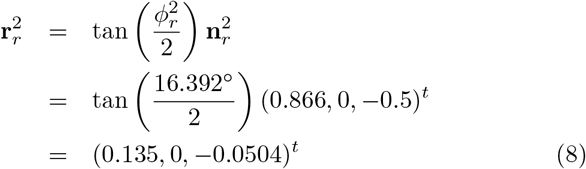

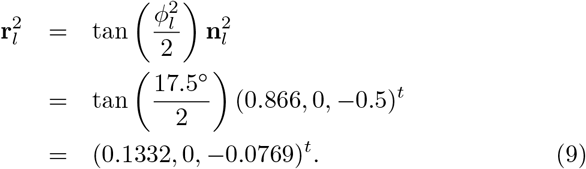

**Figure 5:**
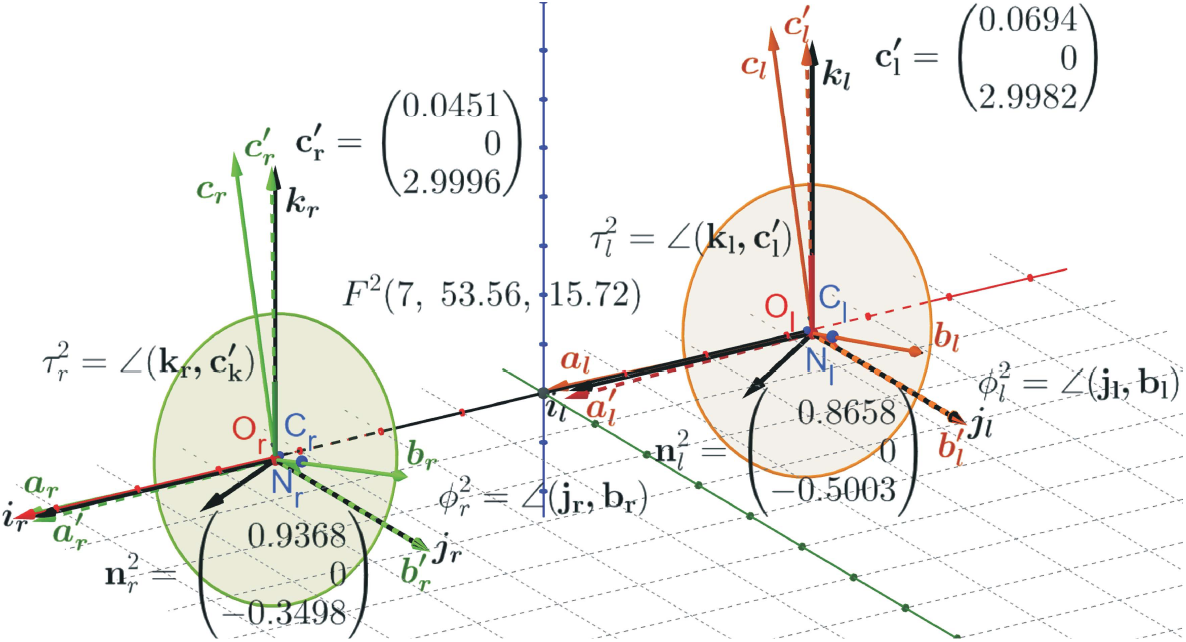
For the fixation *F* ^2^(7, 53.56, 15.72, the frames’ angles of rotations around the axes 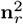 and 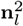 are 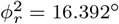 and 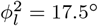, respectively for the right and left AE. Similarly, the ocular torsions are 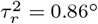 and 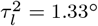.

Then, frames rotations that align vector **b**_*r*_ for *F* ^1^(12, 33.56, 5.72) with with vector **b**_*r*_ for *F* ^2^(7, 53.56, 15.72) and similarly for vectors **b**_*l*_s are calculated as follows. Let these Rodrigues’ vectors be denoted by 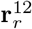 and 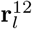 for the right and left eye, respectively. Thus, using the properties of Rodrigues’ vectors discussed above, I can first write **r**^12^ ° **r**^1^ = **r**^2^ so that **r**^12^ = **r**^2^ ° −**r**^1^ and then, using Eq (5), I obtain

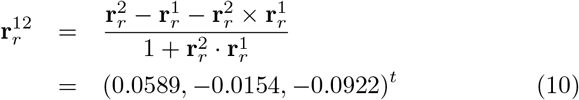

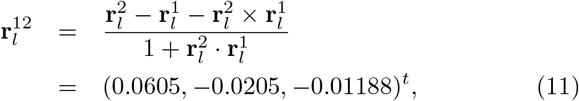

so that from Eqs (2), (10) and (11), I have

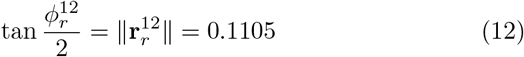

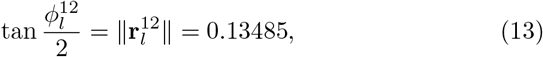

which gives the corresponding angle values 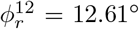 and 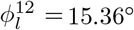 for the right and left eye, respectively.

Finally, these Rodrigues’ vectors can be written as

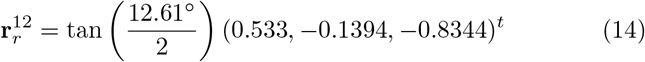

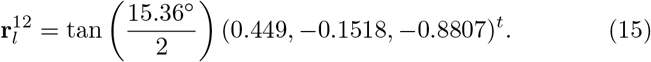

The ocular torsions when the eyes fixation changes from *F* ^1^ to *F* ^2^ is calculated as 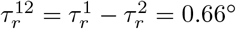 and 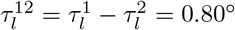 using the values given in Figures 4 and 5.

### 5.3 The Binocular Half-Angle Rule

I conclude this section with the half-angle rule. It describes the rotation of the eye between tertiary configurations first discussed by Helmholtz with the connection to Listing’s law.

The half-angle rule was attributed to the non-commutativity of 3D rotations, see Martinez-Trujillo [2005]. It was first proved in Cannata and Maggiali [2008] using quaternion representation of rotations. Also, its alternative proof was given by more general nonlinear dynamics methods along with an extensive discussion in Novelia and O’Reilly [2015]. These proofs were only given for one eye. Here, the half-angle rule is demonstrated in *GeoGebra* simulations for binocular fixations. However, the formulation may feel subjective because it does not involve time.

The simulation shown in Figure 6 demonstrates that Rodrigues’ vectors 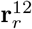 and 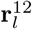 are almost perpendicular to the corresponding bisectors *b*_*r*_ and *b*_*l*_ in the greyish plane and the brownish plane, respectively. These planes contain 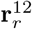 and the perpendicular line *l*_*r*_ to the optical center of ERP image plane for the right eye (*T*_*r*_ axis in Figure 1 for the right eye) and 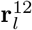 and the perpendicular line *l*_*l*_ to the optical center of ERP image plane for the left eye. Then, for each eye, the visual line of *F* ^1^ is rotated about the intersection line of the corresponding plane introduced above and the frontal plane of ERP—the lines *m*_*r*_ for the right eye and *m*_*l*_ for the left eye. The resulting lines are shown with points 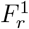 and 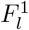. The bisector lines *b*_*r*_ and *b*_*l*_ are between these two sets of lines. It resolves the eccentricity of the fixation *F* ^1^ as it is discussed above.

**Figure 6:**
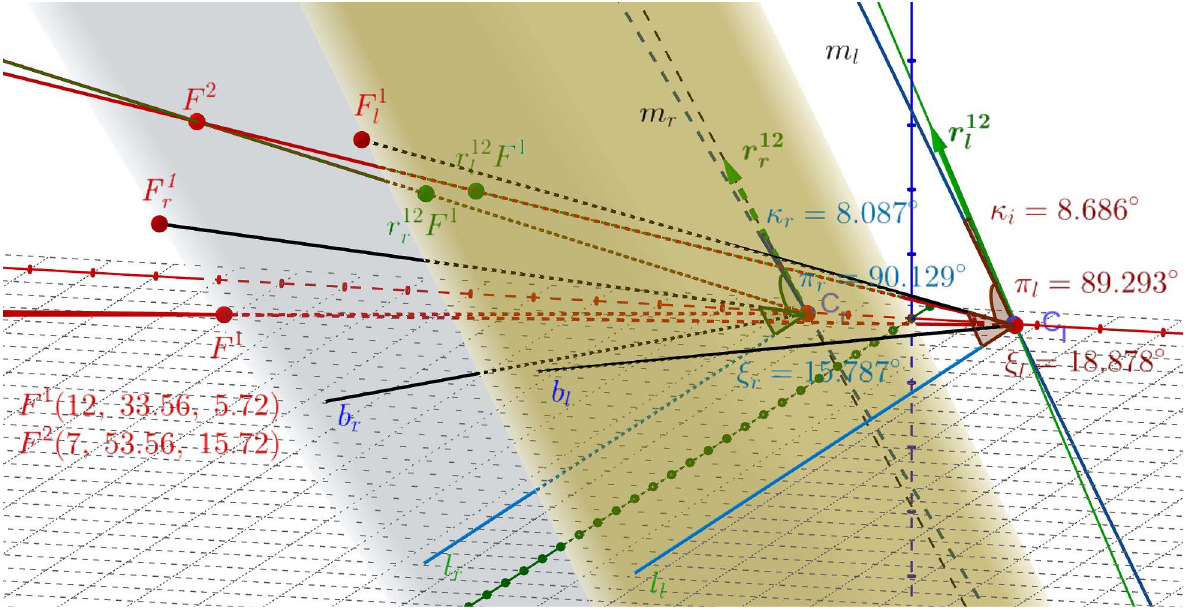
The construction of the binocular half-angle rule. The angles *κ*_*r*_ = 8.1^*°*^ and *κ*_*l*_ = 8.7^*°*^ approximate the half-angle of *ξ*_*r*_ = 15.8^*°*^ and *ξ*_*l*_ = 18.9^*°*^ for the right and the left eye, respectively. The half-angle rule, which is approximate because of the eyes’ misaligned optical components, is demonstrated by the angles *π*_*r*_ and *π*_*l*_ that are very close to right angles. See the discussion in the text for the construction of these angles.

This half-angle rule (when the eye’s misalignment is disregarded) is a corollary of Euler’s rotation theorem for the ERP posture, an appropriate substitution for the primary plane underlying Listing’s law. The real significance of this rule is the fact that it is binocular: it is applied to each eye and transforms the bifoveal fixations from *F* ^1^(12, 33.56, 5.72) to *F* ^2^(7, 53.56, 15.72) by rotating with Rodrigues’ vectors (14) and (15). However, the small discrepancy mentioned above in the text and in the caption of Figure 6 results from the eye’s misaligned optical components. Importantly, the nonzero vergence coefficients that define the tilts of the initial Listing plane for each eye in the binocular extensions L2 that are controversial are not needed in the approach used here.

## 6 Conclusions

In healthy human eyes, the fovea is displaced from the posterior pole, and the lens is tilted away from the optical axis. It is well known that the lens’ horizontal tilt partially compensates for the optical aberrations resulting from the fovea’s misalignment and the asphericity of the cornea. The recent study in Turski [2023] established the geometric theory of horizontal retinal correspondence’s spatial coordinates (iso-disparity curves) in the binocular system with the asymmetric eye (AE) model that comprises horizontal misalignment of optical components. That study revealed that the distribution of the iso-disparity lines is invariant to the AE horizontal misalignment of the lens. It demonstrated for the first time that misaligned optical components cause the observed asymmetry of the retinal correspondence in human eyes.

This work uses the 3D AE model, which comprises horizontal and vertical misalignments of the eye’s optical components, to discuss Listing’s law in oculomotor vision.

Ocular torsion for the 3D AE binocular posture is geometrically defined using Euler’s rotation theorem, which closely approximates the corneal rolling eye movement and is simulated in *GeoGebra* in the framework of Rodrigues’ vector. It provides a consistent binocular formulation of Listing’s law for the first time. In particular, it replaces the Listing plane defined *ad hoc* for a single eye and extends to “binocular” eye movement with parallel gazes with the eye’s resting posture or ERP. The ERP is the binocular posture that, for the average asymmetry angles of the AE, numerically corresponds to the eyes’ resting vergence posture, in which the eye muscles’ natural tonus resting position serves as a zero-reference level for convergence effort Ebenholtz [2001]. The binocular ERP provides a new definition for Listing’s law eye’s primary position—gaze line perpendicular to the Listing plane containing rotation vectors of eye movement executed from the primary position. Note that the eye’s primary position neurophysiological significance has remained elusive despite its theoretical importance in the oculomotor control Hess and Thomassen [2014].

The simulation results of Section 5, which were obtained for tertiary postures when the fixation changed from *F* ^1^(12, 33.56, 5.72) to *F* ^2^(7, 53.56, 15.72), allow us to consider two different descriptions of a sequence of eyes’ binocular posture execution in a stationary upright head. In the first description, the change between the two consecutive postures is related to the preceding and proceeding postures,

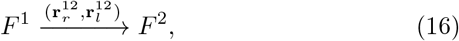

where 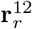 and 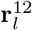 are Rodrigues’ vectors (8) and (9). This first execution in Rodrigues’ vector framework produced the binocular half-angle rule between two tertiary postures in the simulation shown in Figure 6.

On the other hand, in the second description, changes between any two consecutive postures are related to the ERP,

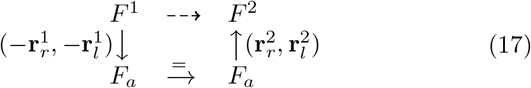

where 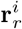 and 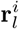 are Rodrigues’ vectors (6) and (7) for *i* = 1 and (8) and (9) for *i* = 2. This is a commutative diagram in which the dashed arrow indicates the composition of transformations given by continuous arrows.

The second description of the consecutive eye binocular posture shifts relative to the ERP is more straightforward and may show how the visual system works. It seems to be supported by experimental observations. In Jampel and Shi [1992], the experimental measurements supported by photographic and video analysis demonstrated that the anatomically determined primary position is a natural constant position to which the eyes are automatically reset from any displacement of the visual line. Further, the evidence indicated an active neurologic basis for the primary position.

The half-angle rule is usually claimed to be satisfied during saccadic and smooth pursuit eye movements [Tweed and Vilis, 1990, Tweed et al., 1992]. However, findings in Thurtell et al. [2011, 2012] indicate that the half-angle rule is not perfectly satisfied during saccades or pursuit in humans.

The theoretical study and simulations presented here, which include the common misalignment of the eye’s optical components, clearly show that the half-angle rule cannot be perfectly satisfied in the human population. However, it is satisfied approximately, and how this approximation depends on the eye’s asymmetry angular parameters should be carefully studied.

Finally, the results of this study suggest that the use of Listing’s law in controlling 3D eye movements, clinical implications for strabismus, and optimal management of this disorder Wong et al. [2002], Wong [2004] should be updated.

## Availability of data and materials

The datasets generated in *GeoGebra* during the current study are available from the corresponding author upon reasonable request.

## References

Y. Chang, H. M. Wu, and Y. F. Lin. The axial misalignment between ocular lens and cornea observed by mri (i) at fixed accommodative state. Vision Res., 47:71–84, 2007. doi: 10.1016/j.visres.2006.09.018.

A. de Castro, P. Rosale, and S. Marcos. Tilt and decentration of intraocular lenses in vivo from purkinje and scheimpflug imaging: validation study. J Cataract Refract Surg., 33:418–429, 2007.

F. Schaeffel and H. Kaymak. New techniques to measure lens tilt, decentration and longitudinal chromatic aberration in phakic and pseudophakic eyes. Nova Acta Leopoldina, 111:127–136, 2010. doi: 10.1167/iovs.07-1022.

G. K. Aguirre. A model of the entrance pupil of the human eye. Sci. Rep., 9(1):9360:1–10, 2019. doi: 10.1038/s41598-019-45827-3.

L. Wang, R. Guimaraes de Souza, M. P. Weikert, and et al. Evaluation of crystalline lens and intraocular lens tilt using a swept-source optical coherence tomography biometer. J Cataract Refract Surg., 45:35–40, 2019. doi: 10.1167/iovs.07-1022.

J. T. Holladay. Quality of Vision: Essential Optics for the Cataract and Refractive Surgeon. SLACK Inc., 2007.

J. Tabernero, A. Benito, E. Alcón, and et al. Mechanism of compensation of aberrations in the human eye. Progress in Brain Research, 24:3274–3283, 2007.

W.N. Charman and D. A. Atchison. Decentred optical axes and aberrations along principal visual field meridians. Vision Research, 49:1869–1876, 2009. doi: 10.1016/j.visres.2009.04.024.

P. Artal. Optics of the eyes and its impact in vision. Adv. Opt. Photon, 6:340–367, 2014. doi: 10.1364/AOP.6.000340.

T. Liu and L. N. Thibos. Variation of axial and oblique astigmatism with accommodation across the visual field. Journal of Vision, 17(3):24:1–23, 2017. doi: 10.1167/17.3.24.

J. Turski. Binocular system with asymmetric eyes. J. Opt. Soc. Am. A, 35:1180–1191, 2018. doi: 10.1364/JOSAA.35.001180.

J. Turski. A geometric theory integrating human binocular vision with eye movement. Front. Neurosci., 14:555965:1–17, 2020. doi: 10.3389/fnins.2020.555965.

J. Turski. Riemannian geometries of visual space: Variable curvature and horizon. Mathematical Methods in the Applied Sciences, 46:9298–9324, 2023. doi: 10.1002/mma.9054.

J. Turski. Geometric study in binocular vision of healthy human eye’s misaligned optical components. bioRxiv, pages 1–32, 2024. doi: 10.1101/2024.06.01.595622.

C. Wheatstone. On some remarkable and hitherto unobserved phenomena of binocular vision. Philos. Trans. R. Soc., 128:371–394, 1838.

B. Julesz. Foundation of Cyclopean Perception. The University of Chicago Press, 1971.

S. M. Ebenholtz. Oculomotor Systems and Perception. Cambridge U. Press, 2001. doi: 10.1017/CBO9780511529795.

B. J. M. Hess and J. S. Thomassen. Kinematics of visually-guided eye movements. PLoS ONE, 9:e95234.1–16, 2014. doi: 10.1371/journal.pone.0095234.

T. Shipley and S. Rawlings. The nonius horopter-i. history and theory. Vis. Res., 10:1225–1262, 1970. doi: 10.1016/0042-6989(70)90039-8.

G. Amigo. A vertical horopter. Optica Acta: International Journal of Optics, 21(4):277–292, 1974.

J. Siderov, R. S. Harwerth, and H. E. Bedell. Stereopsis, cyclovergence and the backward tilt of the vertical horopter. Vision Research, 39:1347–1357, 1999.

H.L.F. Helmholtz. Physiological Optics (Southhall, J.P.C. translation). Optical Society of America, 1867/1925.

S. R. Barry. Beyond the critical period. acquiring stereopsis in adulthood. In Plasticity in Sensory Systems, pages 175–195. Cambridge U. Press, Cambridge, 2013. doi: 10.1017/CBO9781139136907.010.

D. Tweed, W. Cadera, and T. Vilis. Computing three-dimensional eye position quaternions and eye velocity from search coil signals. Vision Research, 30:97–110, 1990.

D. Tweed and T. Vilis. Geometric relation of eye position and velocity vectors during saccades. Vision Research, 30:111–127, 1990.

T. Haslwanter. Mathematics of three-dimensional eye rotations. Vision Research, 35:1727–1339, 1995.

A. Novelia and O.M. O’Reilly. On the dynamics of the eye: geodesics on a configuration manifold, motions of the gaze direction and helmholtz’s theorem. Nonlinear Dyn., 80:1303–1327, 2015. doi: 10.1007/s11071-015-1945-0.

D. Mok, A. Ro, W. Cadera, and et al. Rotation of listing’s plane during vergence. Vision Research, 32:2055–2064, 1992.

A. H. Minken and J. A. M. van Gisbergen. A three-dimensional analysis of vergence movements at various levels of elevation. Experimental Brain Research, 101:331–345, 1994. doi: 10.1007/BF00228754.

L. J. Van Rijn and A. V. Van der Berg. Binocular eye orientation during fixations: Listing’s law extended to include eye vergence. Vision Research, 33:691–708, 1993.

D. Tweed. Visual-motor optimization in binocular control. Vision Research, 37:1939–1951, 1997.

E. Piña. Rotations with rodrigues’ vector. EUROPEAN JOURNAL OF PHYSICS, 32:171–1178, 2011.

J. J. Gray. Olinde rodrigues’ paper of 1840 on transformation groups. Arch. Hist. Exact Sci., 21:375–384, 1980.

J. C. Martinez-Trujillo. Noncommutativity of eye rotations and the half-angle rule. Neuron, 47:171–173, 2005. doi: 10.1016/j.neuron.2005.07.004.

G. Cannata and M. Maggiali. Models for the design of bioinspired robot eyes. IEEE Transactions on Robotics, 24:27–44, 2008. doi: 10.1109/TRO.2007.906270.

R. S. Jampel and D. X. Shi. The primary position of the eyes, the resetting saccades, and the transverse visual head plane. Investigative Ophthalmology and Visual Science, 33:2501–2510, 1992.

D. Tweed, M. Fetter, S. Andreadaki, E. Koenig, and J. Dichgans. Three-dimensional properties of human pursuit eye movements. Vision Research, 32:1225–1238, 1992.

M. J. Thurtell, A. C. Joshi, R. J. Leight, and M. F. Walker. Three-dimensional kinematics of saccadic eye movements in humans: is the “half-angle rule” obeyed? Ann N.Y. Acad Sci., 1233:34–40, 2011.

M. J. Thurtell, A. C. Joshi, and M. F. Walker. Three-dimensional kinematics of saccadic and pursuit eye movements in humans: Relationship between donders’ and listing’s laws. Vision Research, 60:7–15, 2012.

A. M. F. Wong, J. A. Sharpe, and D. Tweed. Adaptive neural mechanism for listing’s law revealed in patients with fourth nerve palsy. Invest. Ophthalmol. Vis. Sci., 43:1796–803, 2002.

A. M. F. Wong. Listing’s law: Clinical significance and implications for neural control. Survey of Ophthalmology, 49:563–575, 2004. doi: 10.1016/j.survophthal.2004.08.002.

